# Structural basis for Rab6 activation by the Ric1-Rgp1 complex

**DOI:** 10.1101/2024.05.06.592747

**Authors:** J. Ryan Feathers, Ryan C. Vignogna, J. Christopher Fromme

## Abstract

Rab GTPases act as molecular switches to regulate organelle homeostasis and membrane trafficking. Rab6 plays a central role in regulating cargo flux through the Golgi and is activated via nucleotide exchange by the Ric1-Rgp1 protein complex. Ric1-Rgp1 is conserved throughout eukaryotes but the structural and mechanistic basis for its function has not been established. Here we report the cryoEM structure of a Ric1-Rgp1-Rab6 complex representing a key intermediate of the nucleotide exchange reaction. This structure reveals the overall architecture of the complex and enabled us to identify interactions critical for proper recognition and activation of Rab6 on the Golgi membrane surface. Ric1-Rgp1 interacts with the nucleotide-binding domain of Rab6 using an uncharacterized helical domain, which we establish as a novel RabGEF domain by identifying residues required for Rab6 nucleotide exchange. Unexpectedly, the complex uses an arrestin fold to interact with the Rab6 hypervariable domain, indicating that interactions with the unstructured C-terminal regions of Rab GTPases may be a common specificity mechanism used by their activators. Collectively, our findings provide a detailed mechanistic understanding of regulated Rab6 activation at the Golgi.

## INTRODUCTION

The Golgi complex is the central hub for protein modification and sorting in eukaryotic cells. Membrane trafficking at the Golgi is regulated by Rab GTPases that function as molecular switches on the cytoplasmic surface of Golgi compartments by recruiting effector machinery to promote vesicle budding, docking, and fusion^1–3^. Rab GTPases are activated by guanine nucleotide exchange factors (GEFs), which are necessary and sufficient for determining where and when Rab proteins are active^3^.

Once activated, Rab proteins are bound to their target membranes via hydrophobic prenyl modifications of an unstructured C-terminal region referred to as the hypervariable domain (HVD)^4^. The HVD is so named to denote how its sequence varies between different Rab proteins, belying the fact that these HVDs tend to be conserved among the same Rab paralogs across different species.

Rab6 is a central regulator of Golgi trafficking and cellular homeostasis. Rab6 is known to regulate endosome-to-Golgi trafficking and autophagy in both budding yeast and metazoans^5–13^. In metazoans Rab6 has also been implicated in secretory traffic from the *trans*-Golgi network^14^. Loss of Rab6 function is embryonic lethal in mice^15^ and renders budding yeast temperature sensitive^16^.

The Ric1-Rgp1 protein complex is a conserved guanine nucleotide exchange factor (GEF) that is known to activate Rab6 in budding yeast^8^ and humans^17^. Dissection of the human complex determined that both the RGP1 subunit and a C-terminal region of the RIC1 subunit interact with human RAB6A, while the middle region of RIC1 interacts with RGP1. In addition, Ric1-Rgp1 was identified as a potential effector of Rab33^17^.

It remains unresolved how Ric1-Rgp1 specifically recognizes Rab6 and the mechanism by which it catalyzes Rab6 nucleotide exchange. Furthermore, little is known about the structure and domain organization of Ric1-Rgp1. Other Rabs, including Rab1, Rab7, and Rab11, are also activated by multi-subunit GEFs^18–21^, but Ric1-Rgp1 does not share any sequence similarity with these other GEF complexes.

To understand how Ric1-Rgp1 activates Rab6 at the Golgi, we used nucleotide depletion to trap a key intermediate step of the nucleotide exchange reaction and then determined its structure by cryo-electron microscopy (cryoEM). The structure reveals the overall architecture of the complex, which is distinct from other known RabGEFs, and has enabled us to identify interactions that are critical for proper recognition and activation of Rab6 on the membrane surface. Ric1 is composed of two β-propellers and a C-terminal α;-solenoid domain that remodels the Rab6 nucleotide binding pocket in order to facilitate nucleotide exchange. Rgp1 posseses an arrestin domain that binds directly to a portion of the otherwise unstructured HVD of Rab6. Accordingly, we find that the Rab6 HVD is important for robust activation of Rab6 *in vivo*. We identify a predicted amphipathic helix within the Rgp1 subunit that is important for Golgi localization. Our findings provide a detailed understanding of the molecular mechanisms underlying regulated Rab6 activation by the Ric1-Rgp1 complex and identify conserved structural elements in both subunits that are required for proper localization, function, and specificity.

## RESULTS

### Architecture of the Ric1-Rgp1-Rab6 activation intermediate complex

To enable structural studies of the Ric1-Rgp1 complex and its interaction with Rab6, we purified the endogenous Ric1-Rgp1 complex from *S. cerevisiae* using C-terminal affinity tags. We then prepared a stable complex with full-length Rab6 encoded by the *S. cerevisiae YPT6* gene by depleting its bound guanine nucleotide. We determined the structure of the resulting nucleotide exchange intermediate complex using cryoEM (Fig. 1a, Table 1, and Supplementary Figs. S1-S2). The global resolution of the cryoEM map is 3.3 Å (using the 0.143 FSC cutoff), enabling confident model building (Fig. 1b-d). We used the resulting model to assign structural domains to the primary structure of both subunits (Fig. 1e).

**Figure 1:**
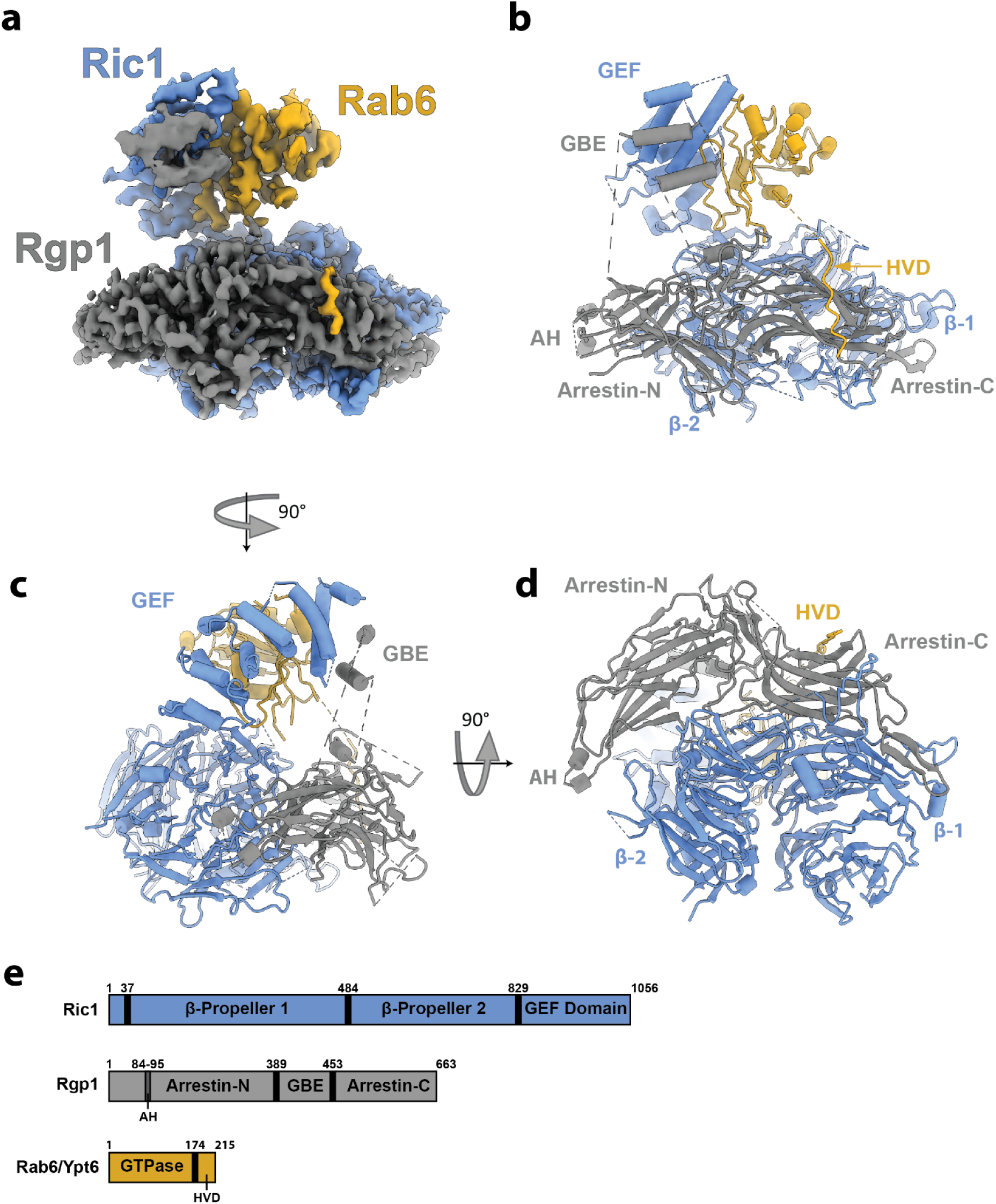
Architecture of the Ric1-Rgp1-Rab6 activation intermediate complex. a) CryoEM reconstruction of the Ric1-Rgp1 complex ‘caught in the act’ of performing nucleotide exchange on Rab6. b) Atomic model of the complex built using the cryoEM density, shown in cartoon format, and highlighting several structural elements including some newly defined in this work (see panel e). c) Same as in (b), from a different perspective. d) Same as in (b) and (c), from a different perspective. e) Structural elements of the three proteins in the complex. Newly established elements are the GEF domain, the GBE (GEF binding element); and an amphipathic helix (AH).

**Table 1:**
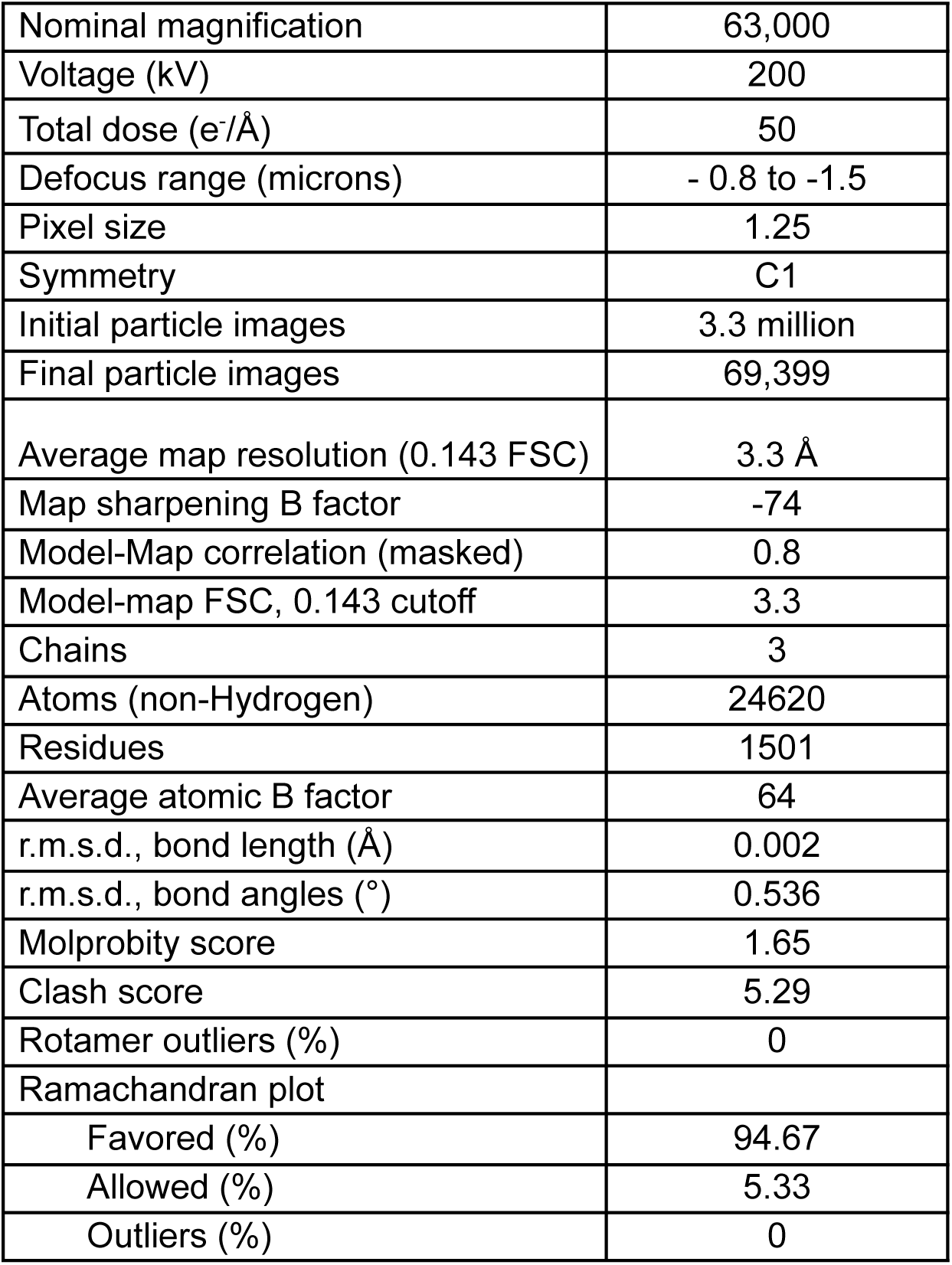
CryoEM data and model statistics.

The structure reveals that the Ric1-Rgp1 complex adopts a two-lobed structure. The larger lobe consists of the bulk of both proteins, including the two β-propeller domains of Ric1 that adopt a clamshell arrangement. Rgp1 adopts an arrestin fold that wraps around the Ric1 β-propellers, burying a large (∼3800 Å^2^) surface area. The arrestin-C subdomain forms a groove that interacts with the HVD of Rab6 (Fig. 1a,b,d) and will be discussed further below.

The small lobe of the Ric1-Rgp1 complex contains the C-terminal portion of Ric1, which adopts an α;-solenoid fold. This domain of Ric1 is bound to the nucleotide-binding domain of Rab6 (Fig 1a-d), consistent with the previous finding that the C-terminal portion of human RIC1 was sufficient for interaction with RAB6A^17^. We refer to this C-terminal α;-solenoid domain of Ric1 as the GEF domain (Fig. 1e) as we have biochemically established its nucleotide exchange activity as described further below. The small lobe also contains two α;-helices of Rgp1 that serve to cap the C-terminus of Ric1, burying ∼600 Å^2^ surface area. These two conserved α;-helices are embedded within a flexible linker region that lies between the Rgp1 arrestin-N and arrestin-C subdomains in the primary sequence and extends from the large lobe. We refer to these two α;-helices of Rgp1 as the GEF domain binding element (GBE) (Fig 1b,e).

### The basis for Ric1-Rgp1 catalysis of Rab6 nucleotide exchange

Rab GTPases adopt distinct conformations in their inactive and active states. In the GTP-bound active state, ‘switch’ regions adopt a conformation that functions as a binding site for effectors^22^. In the nucleotide-free state captured in the cryoEM structure, Ric1 wraps around the surface of Rab6 and makes extensive contacts with the switch regions of the GTPase, burying ∼1350 Å^2^ surface area at the GEF domain interface (Figs. 1a-c, 2a). While most of the interaction involves the GEF domain, the second β-propeller of Ric1 also contacts Rab6, creating a smaller interface of ∼350 Å^2^.

The ‘switch I’ region observed in the structure of the nucleotide-free complex adopts a conformation that is dramatically different from known structures of Rab6 in the active and inactive states^23,24^. The interaction with the GEF domain has peeled switch I away from the guanine nucleotide binding-site, with several residues involved in nucleotide binding displaced by ∼25 Å compared to their positions in the GDP-bound state (Fig 2b, Video S1). This displacement effectively eliminates the nucleotide-binding pocket, explaining how Ric1-Rgp1 releases the bound nucleotide. Accordingly, residues comprising the P-loop and a portion of the switch II region are not visible in the cryoEM density.

**Figure 2:**
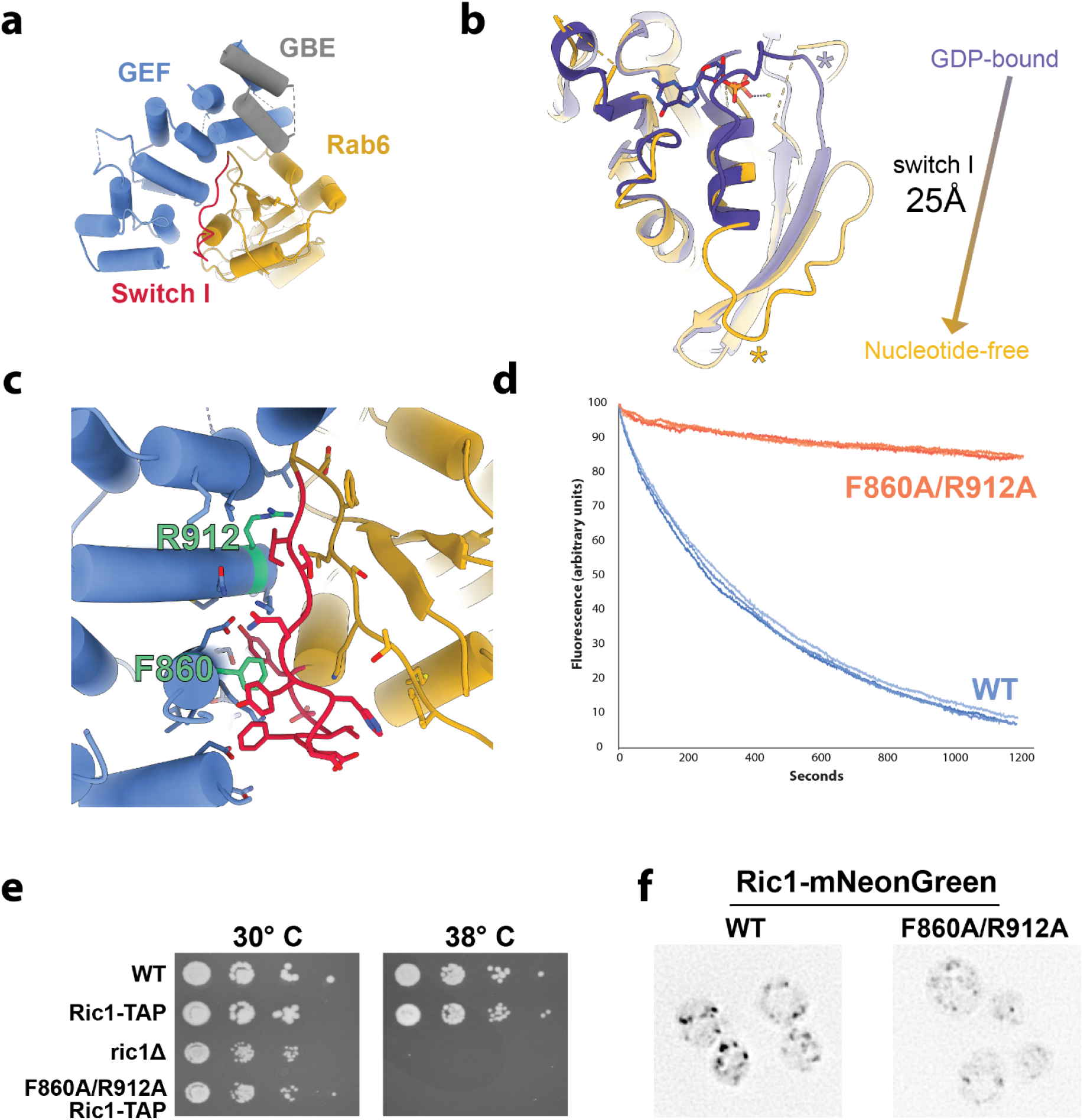
Structural basis for Rab6 activation by nucleotide exchange. a) View of the interface between the Ric1 GEF domain (blue) and Rab6 (gold), with the Rab6 ‘switch I’ element (residues 32-44) colored red. The GBE of Rgp1 is colored gray. b) Overlay of the crystal structure of Rab6 in its GDP-bound state (colored purple, PDB: 1D5C)^23^, with the nucleotide-free state from the Ric1-Rgp1-bound cryoEM structure (colored gold). Equivalent positions in primary sequence are marked with asterisks. c) Close-up view of the Rab6-GEF domain interface, highlighting the Ric1 residues subjected to mutational analysis. d) Results from an *in vitro* GEF assay using purified Ric1-Rgp1, prenylated-Rab6-GDI complex, and liposomes. The change in fluorescence is due to exchange of the Rab6-bound mant-GDP for GTP, representing activation of Rab6. n = 3 traces are shown for each condition (WT or F860A/R912A Ric1-Rgp1 complex). e) Complementation test (cells lacking Ric1-Rgp1 function are temperature sensitive). f). Comparison of the localization patterns of WT and F860A/R912A Ric1-mNeonGreen.

At the core of the interface between Ric1 and Rab6, residues Phe860 and Arg912 of the GEF domain make direct contact with the switch I region of Rab6 (Fig. 2c). To test whether these Ric1 residues are required for nucleotide exchange we utilized a physiological *in vitro* RabGEF activity assay^25,26^. This assay utilizes a fluorescent GDP analog to measure the kinetics of nucleotide exchange of Rab6 in the presence of synthetic liposome membranes with a lipid composition that mimics that of the Golgi^27^. *In vivo*, Rab6 is geranylgeranylated on both of its C-terminal cysteines and inactive Rab6-GDP is kept soluble in the cytosol via masking of these hydrophobic prenyl groups by the chaperone protein GDI^28,29^. We therefore used a prenylated Rab6-GDI complex prepared by enzymatic synthesis (Supplementary Fig. S1c) as the Rab substrate for the *in vitro* GEF reactions. While such physiological GEF assays have been performed for other RabGEF-Rab pairs^25,30^, to our knowledge this is the first time such a physiological GEF assay has been established for Rab6.

Using this assay, we found that the purified wild-type Ric1-Rgp1 complex exhibited robust GEF activity on the prenylated Rab6-GDI complex in the presence of liposomes (Figs. 2d, S1d). In contrast, the Ric1 F860A/R912A mutant complex was catalytically dead, indicating Ric1 residues Phe860 and Arg912 together play a critical role in Rab6 nucleotide exchange by the Ric1-Rgp1 complex.

Rab6 activation is required for growth of *S. cerevisiae* at high temperatures, consequently cells lacking Rab6, Ric1, or Rgp1 are temperature sensitive^16^. To test the importance of Phe860 and Arg912 *in vivo* we examined the growth of yeast cells harboring mutant Ric1 constructs. We found that the Ric1 F860A/R912A mutant was also temperature-sensitive (Fig. 2e), indicating this mutant has lost Ric1 function *in vivo*. This mutant was expressed, albeit at reduced levels (Supplementary Fig. S1e). Similar to the wild-type, the mutant protein appeared to localize to punctate Golgi compartments, but these puncta appeared smaller and more numerous compared to the wild-type (Fig. 2f). Loss of Rab6 function is known to result in smaller and more numerous Golgi compartments^31^, so our interpretation of these results is that the F860/R912A mutant Ric1-Rgp1 complex is able to bind to the Golgi membrane but unable to activate Rab6, thus altering Golgi morphology. Taken together, these results indicate that Phe860 and Arg912 of Ric1 together are required for the Rab6 nucleotide exchange function of the Ric1-Rgp1 complex *in vivo*, and confidently establish the C-terminal Ric1 α;-solenoid as a Rab6 GEF domain.

### Comparison of the Ric1-Rgp1 complex to other GEFs

Based on primary sequence, the Ric1 and Rgp1 subunits do not share any obvious homology to any of the other known RabGEFs. Correspondingly, the cryoEM structure indicates that the overall architecture of the complex does not resemble that of any of the structurally characterized RabGEFs. To assess whether the Ric1 GEF domain might share any structural similarity with other RabGEFs, we compared the structure of the Ric1 GEF domain-Rab6 complex to the structures of several other RabGEFs in complex with their substrate Rab GTPases (Fig. 3a). This comparison illustrates the diversity of GEF-domain folds that cells have evolved to destabilize the nucleotide-binding sites of Rab GTPases. The α;-solenoid fold of the Ric1 GEF domain is distinct from other known RabGEF domain structures, and more closely resembles the “Sec7” Arf-GEF domain, which also adopts an α;-solenoid fold but engages with its substrate GTPase in a distinct manner. (Fig. 3b)^44,45^.

**Figure 3:**
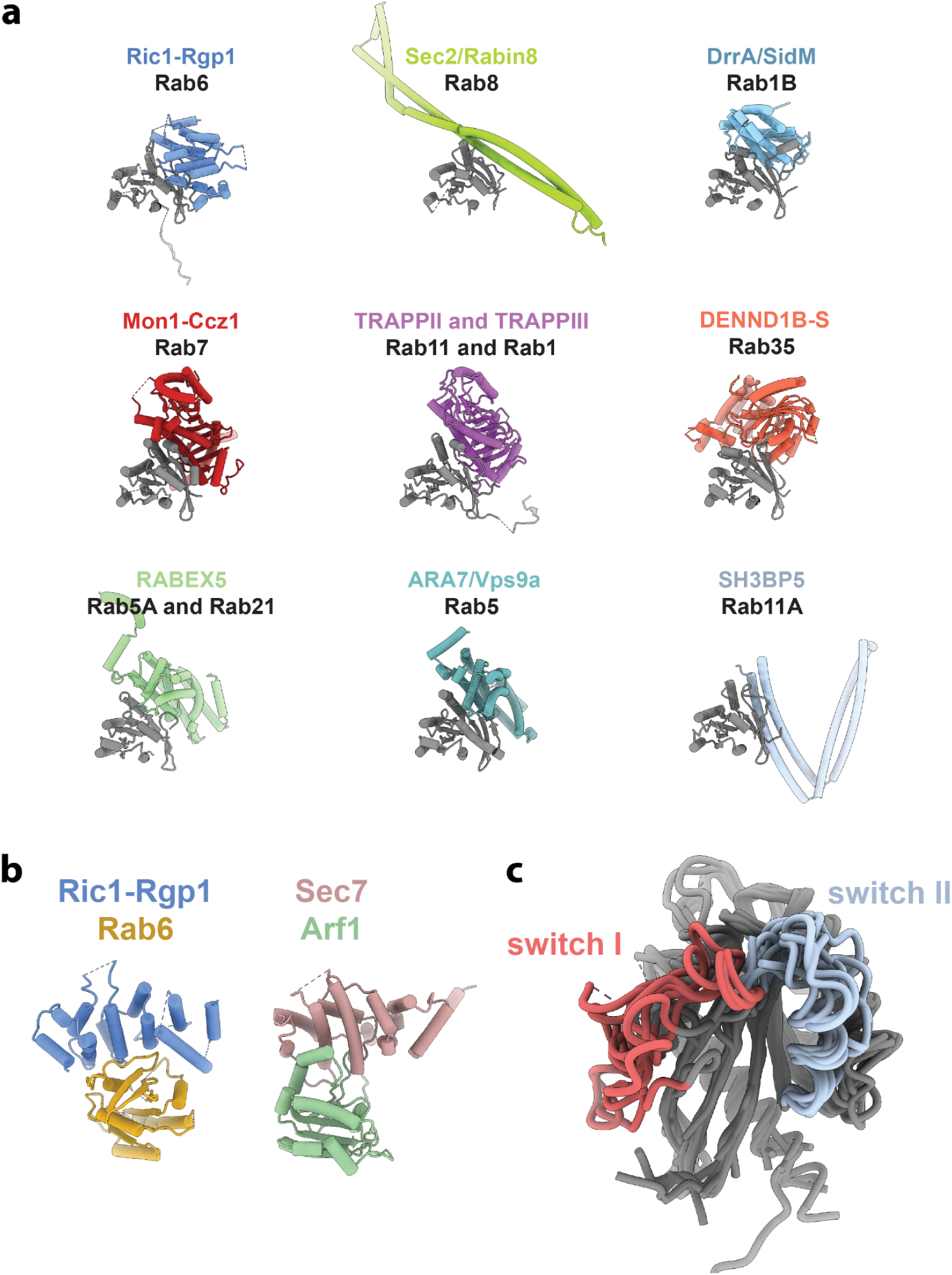
Comparison of Ric1-Rgp1 to other RabGEF domains. a) Comparison of the Ric1 GEF-Rab6 structure to structures of other known RabGEF domain-Rab structures. Note for many of these structures, while the structure of the GEF domain is known, the structure of the intact GEF has not been determined. Each Rab is colored gray and positioned in the same orientation. Ric1-Rgp1 (this work, PDB 9AYR), Sec2/Rabin8^32^, (PDB 2OCY); DrrA/SidM^33^, (PDB 3JZA); Mon1-Ccz1^34^, (PDB 5LDD); TRAPP complexes^35–38^, (PDB 7U05); DENND1B-S^39^, (PDB 3TW8); RABEX-5^40,41^, (PDB 4Q9U); ARA7/Vps9a^42^, (PDB 4G01); SH3BP5^43^, (PDB 6DJL). b) Comparison of the Rab6-bound Ric1 GEF domain to the Arf1-bound Sec7 GEF domain (PDB: 1RE0)^44^. c) Overlay of the nine GEF-bound Rab structures shown in panel (a), with switch I regions colored red and switch II regions colored blue.

The precise conformations adopted by each of the Rabs when bound to its GEF in the nucleotide-free state also varies among the different structures (Fig. 3c), highlighting the distinct mechanisms used by different RabGEFs to perturb nucleotide binding.

### The Rab6 HVD is important for activation at the Golgi

In addition to the interaction between the Ric1 GEF domain and the Rab6 nucleotide-binding domain, the arrestin-C subdomain of the Rgp1 subunit interacts with a portion of the Rab6 HVD (Fig. 4a-c), consistent with the finding that the human RGP1 subunit was able to interact with human RAB6A *in vitro*^17^. Residues 195-204 of the Rab6 HVD are visible in the cryoEM density, bound to a pocket on the surface of Rgp1 and structured as a β-strand integrated on the edge of a β-sheet within the β-sandwich fold of the arrestin-C subdomain (Fig. 4b,c).

**Figure 4:**
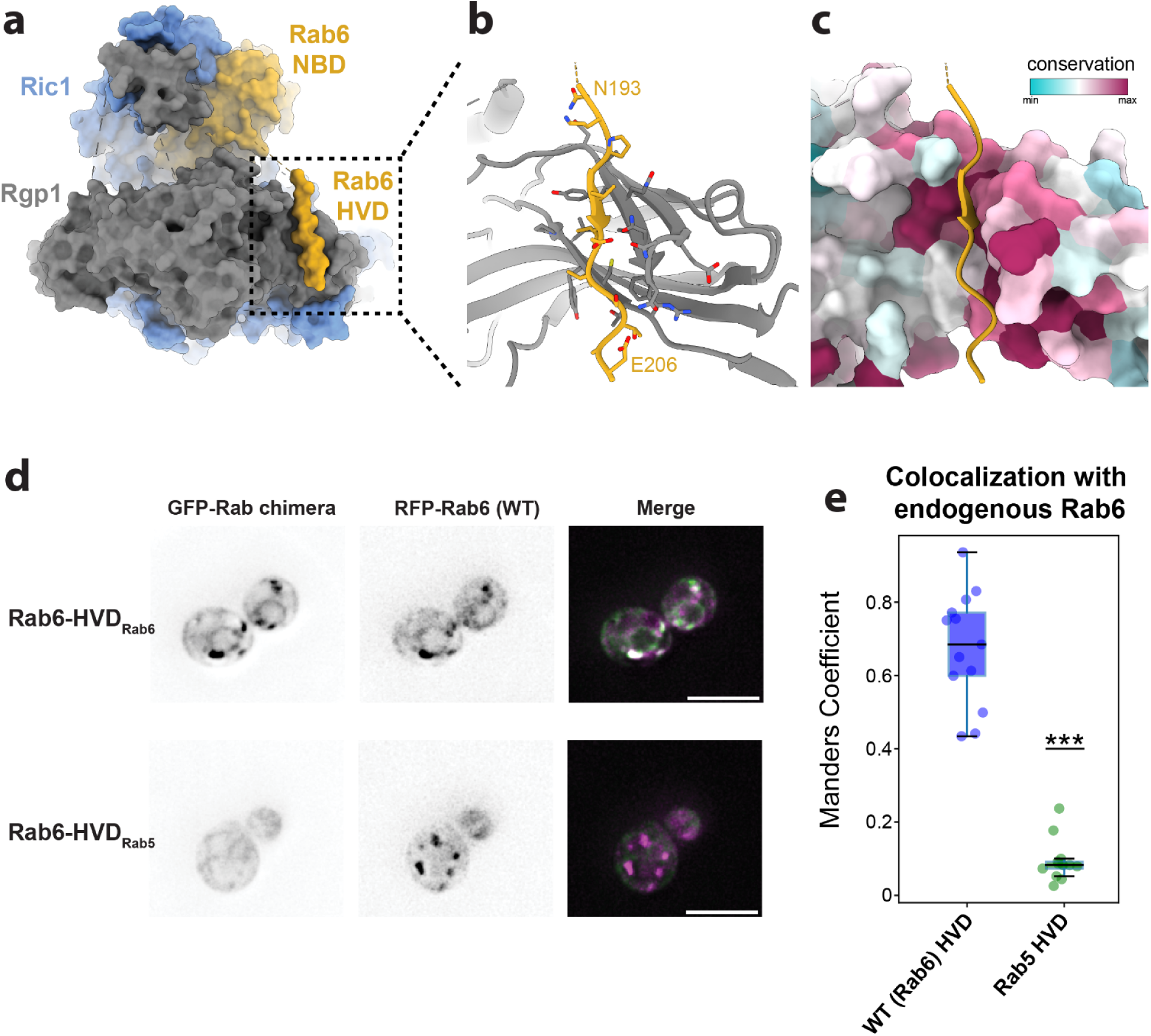
The Ric1-Rgp1 complex binds to the HVD of Rab6. a) Surface view of the Ric1-Rgp1-Rab6 atomic model. b) Close-up view of the portion of the Rab6 HVD bound to the arrestin-C subdomain of Rgp1. The HVD adopts a β-strand conformation to integrate into the β-sheet on the surface of the arrestin-C subdomain β-sandwich fold. c) Same view as in panel (b), with the surface of Rgp1 colored by sequence conservation. d) Live-cell imaging to measure the colocalization of WT and chimeric GFP-tagged Rab6 constructs with endogenous RFP-Rab6. Scale bar, 5 microns. e) Quantitation of the data in (c), with all individual data points overlaid on box-and-whiskers plots. ***, p=2.6×10^-9^. WT: n=13 images; mutant: n=16 images.

While the sequence of the HVD varies between different Rab GTPases, the HVDs of individual Rabs have regions of sequence that are conserved across different organisms (Supplementary Fig. S3a). The sequence of the Rab6 HVD that is bound to Rgp1 includes the ‘CIM’ motif required for prenylation of Rabs by the geranylgeranyl transferase machinery^46^.

There are two other RabGEFs known to bind directly to the HVDs of their substrates, the TRAPPII and TRAPPIII complexes^26,36,37,47^. The TRAPP complexes use their shared Trs31 subunit to bind to the Rab1 and Rab11 HVDs, also encompassing their CIM motifs, similar to the Rgp1-HVD interaction (Supplementary Fig. S3b).

To test the importance of the HVD sequence for the activation of Rab6 by Ric1-Rgp1 at the Golgi *in vivo*, we generated a chimeric GFP-Rab6 construct in which the Rab6 HVD was replaced by the HVD from Rab5 (encoded by the yeast *VPS21* gene). We then visualized the localization of this construct expressed from a centromeric plasmid in cells also expressing endogenously tagged wild-type RFP-Rab6. Previous studies have found that loss of Rab activation *in vivo* results in diffuse localization and/or ectopic localization of the Rab to the endoplasmic reticulum^48^. The wild-type GFP-Rab6 construct exhibited normal punctate localization and colocalized very well with endogenous RFP-Rab6, indicative of localization to the Golgi complex and therefore robust activation (Fig 4d,e). In contrast, the chimeric GFP-Rab6-HVD_Rab5_ construct was significantly mis-localized, with a localization pattern that primarily resembled that of the endoplasmic reticulum, with only very faint and occasional co-localization with RFP-Rab6 (Fig 4d,e). Therefore the sequence of the Rab6 HVD is important for robust activation of Rab6 *in vivo*, likely by contributing to RabGEF specificity.

To determine the impact of swapping the Rab6 HVD on cell viability we introduced the chimeric Rab construct into *rab6Δ* mutant cells. We observed no growth defects at 38°C (Supplementary Fig. S3c), indicating that the low level of GFP-Rab6-HVD_Rab5_ chimera activation was sufficient to support growth at high temperature. We note that previous studies have documented examples of cells with a dramatic loss of essential Rab activation at the Golgi *in vivo* that were nevertheless viable^49–51^. Interestingly, the chimeric construct also exhibited reduced protein levels, perhaps suggesting increased degradation of the inactive Rab chimera (Supplementary Fig. S3d). Taken together, our results indicate that the identity of the HVD is important for robust Rab6 activation in cells but is not essential for activation of the Rab6 pool to a level that supports growth at high temperature.

### Model for Ric1-Rgp1 orientation on the membrane surface

The position and orientation of the Rab6 HVD bound to Rgp1 indicates the membrane-inserting prenyl moieties covalently linked to the C-terminal cysteine residues of Rab6 are located at the ‘bottom’ of the complex as viewed in Fig. 1a. This suggests that the bottom surface of the Ric1-Rgp1 complex may interact directly with Golgi membrane lipid headgroups. The overall electrostatic potential of this surface appears to be somewhat acidic (Supplementary Fig. S4a), rather than possessing a basic character more typical of many peripheral membrane proteins. However, there are several loops projecting from this surface that are disordered and therefore not included in the electrostatic potential calculation of the cryoEM model. Intriguingly, a projection from this surface of the Rgp1 subunit is visible in the cryoEM density at lower thresholds (Fig. 5a). This projection was also apparent in 2D class averages of the cryoEM particle images (Fig. 5b). Analysis of the AlphaFold^52^ structural prediction of Rgp1 suggests that this projection corresponds to two α;-helices extending from the arrestin-N domain (Fig 5c).

**Figure 5:**
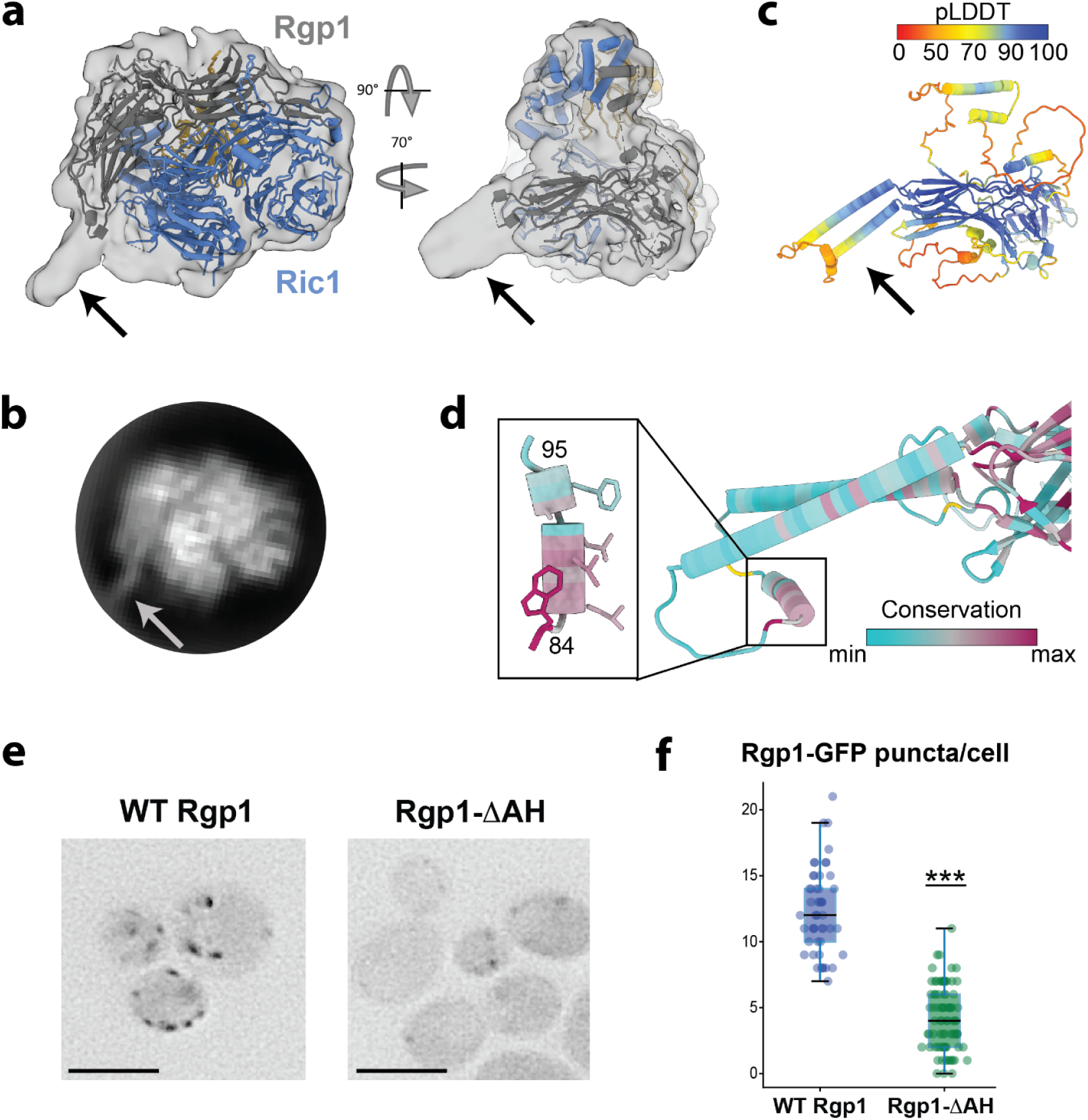
A predicted amphipathic helix in Rgp1 is important for Golgi membrane association. a) CryoEM density map shown at low threshold, together with the atomic model of the Ric1-Rgp1-Rab6 complex. The arrow denotes unmodeled density projecting from the Rgp1 subunit. b) CryoEM 2D class average representing the 2D projection of the complex from the same approximate orientation as the perspective depicted in panel (a). The arrow corresponds to the unmodeled density in the 3D reconstruction. c) AlphaFold predicted structure of the Rgp1 subunit, colored by confidence score. The arrow corresponds to the region of the prediction that is unmodeled in the cryoEM structure due to the low-resolution of that portion of the cryoEM density. d) A conserved amphipathic helix lies at the distal tip of this structural element. e) Live-cell imaging of WT and ΔAH Rgp1-GFP constructs. Scale bar, 5 microns. f) Quantitation of the experiment shown in panel (e) with all individual data points overlaid with box-and-whiskers. ***, p=2.2×10^-16^. WT: n=50 cells; mutant: n=83 cells. See Methods for descriptions of statistics.

The AlphaFold prediction also suggests the presence of a conserved amphipathic helix located at the distal end of these extended helices (Fig. 5d). Given its conserved amphipathic nature, lack of observable density (even at low cryoEM map thresholds), and its flexible connection to the rest of the complex, we viewed this amphipathic helix as a strong candidate for a potential membrane-binding element.

To investigate the functional significance of this amphipathic helix, we generated a mutant version of Rgp1-GFP lacking residues 84-95, which encompass the helix. When introduced into *rgp1Δ* cells, this construct (Rgp1-ΔAH) was significantly mislocalized to the cytoplasm in comparison to the wild-type construct (Fig. 5e,f). Although some punctate localization of the mutant was observed, both the number of puncta and their intensity was significantly diminished. This suggests that the conserved amphipathic helix in Rgp1 is important, but not essential for Golgi localization of the Ric1-Rgp1 complex. The Rgp1-ΔAH construct was able to complement the temperature sensitive phenotype of *rgp1Δ* mutant cells (Supplementary Fig. S4b), and exhibited a somewhat reduced protein level (Supplementary Fig S4c). Although we cannot rule out the possibility that this helix is important for some other aspect of Ric1-Rgp1 GEF function, our interpretation of these results is that the amphipathic helix serves as a membrane binding element, and other factors are also involved in localizing Ric1-Rgp1 to the Golgi.

Based on the locations of the Rab6 C-terminus and the Rgp1 amphipathic helix in the cryoEM structure of the complex, we are able to propose a plausible model for a specific orientation of the Ric1-Rgp1 complex when it is performing Rab6 nucleotide exchange on the membrane surface (Fig. 6).

**Figure 6:**
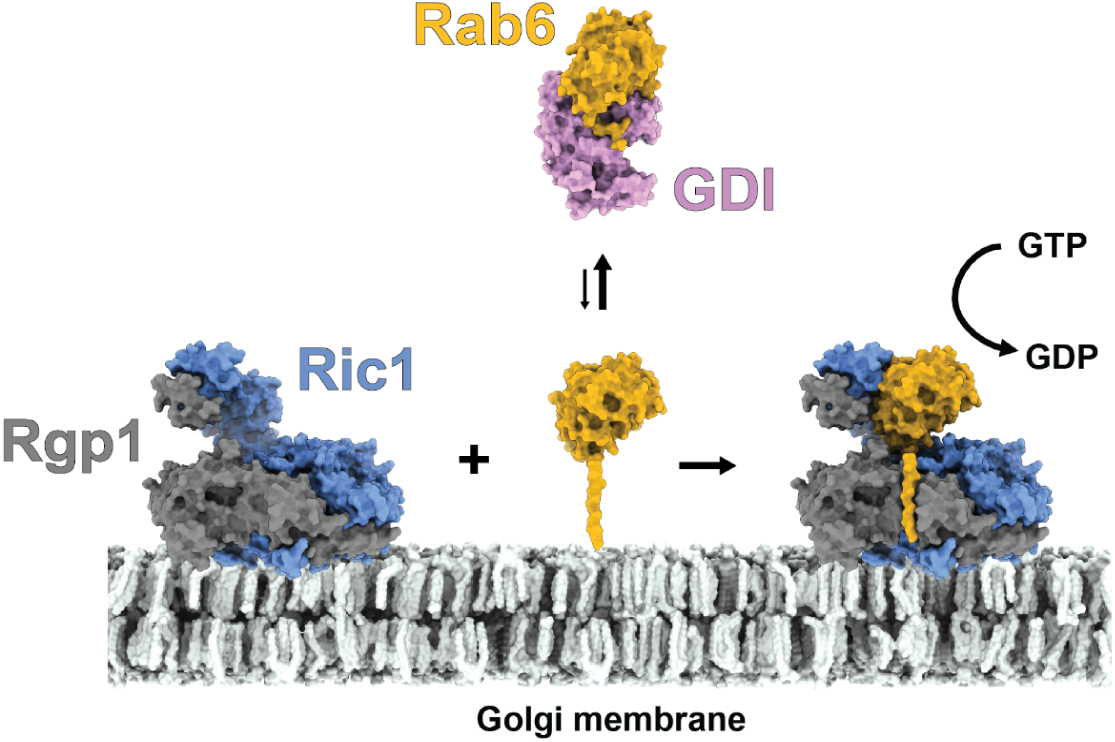
Structural model for Rab6 activation on the Golgi membrane surface. The expected orientation of the Ric1-Rgp1 complex relative to the Golgi membrane surface is shown. GDI must transiently release inactive, GDP-bound Rab6 before it can be activated by the Ric1-Rgp1 complex. The GDI-Rab-GDP complex is depicted using the crystal structure of a GDI-Rab1/Ypt1 complex (PDB: 2BCG)^53^.

## DISCUSSION

Regulated Rab6 activation is essential in eukaryotes for maintaining normal intracellular trafficking and Golgi homeostasis. Rab6 is one of only six Rabs conserved from budding yeast to humans^54–56^. This conservation across distant organisms highlights its importance as a fundamental component of the eukaryotic endomembrane system (Pereira-Leal & Seabra, 2001). Ric1-Rgp1 is known to serve as the GEF for Rab6 but prior to this study its structure and mechanism were unknown. A previous study identified a region at the C-terminus of Ric1 that preferentially binds Rab6-GDP and was able to catalyze nucleotide exchange *in vitro*^17^. In cells however, both Ric1 and Rgp1 are required for Rab6 activation and the nature of the protein-protein interactions between Ric1, Rgp1, and Rab6 has remained unknown.

In this study we have determined the cryoEM structure of the Ric1-Rgp1 complex bound to Rab6 and dissected the mechanisms underlying activation of Rab6 on the Golgi membrane surface. Our cryoEM data reveals the subunit and domain architecture of the Ric1-Rgp1 complex and captures its interaction with Rab6 in the nucleotide-free state.

Structural analysis of the Ric1-Rgp1-Rab6 complex revealed that Ric1 and Rgp1 form a unique architecture that is distinct from other known GEF proteins. We found that Ric1 consists of two β-propellers and an α;-solenoid repeat domain at the C-terminus. A larger region of Ric1 encompassing this C-terminal domain was previously identified to bind Rab6-GDP^17^. The structure presented in this work reveals that a portion of Rgp1, which we term the “GBE”, folds together with the α;-solenoid of Ric1. Using mutational analysis we have now established the role of this structurally distinct RabGEF domain in activation of Rab6 via nucleotide exchange.

We identified an interface formed between the GEF domain and the surface of Rab6 at the switch I loop. We determined that within this interface, residues F860 and R912 of Ric1 together play a critical role in the nucleotide exchange mechanism. Expressing a Ric1 F860A R912A mutant failed to rescue the temperature sensitive growth phenotype of a *ric1Δ* yeast strain. These results are supported by the complete lack of *in vitro* GEF activity exhibited by the corresponding purified mutant protein complex. Taken together, our results indicate the interaction of Ric1-Rgp1 with the Rab6 switch I region destabilizes the nucleotide binding pocket to catalyze guanine nucleotide exchange.

Amphipathic helices are common membrane-binding elements used by regulatory proteins that function at the Golgi. Several Arf and Rab GEFs and GAPs are known to use amphipathic helices to bind to the Golgi membrane^36,57–60^. Our data suggest that the Rgp1 amphipathic helix is involved in Golgi membrane binding, yet it remains unclear how the Ric1-Rgp1 complex is recruited specifically to the surface of the Golgi, rather than another organelle. We observed that removing the Rgp1 amphipathic helix dramatically reduced, but did not completely eliminate, the punctate localization of Rgp1 in cells, implying the presence of an additional interaction with an unknown partner that serves to localize the complex. A previous report identified a direct interaction between the human RIC1 C-terminus and GTP-bound RAB33B^17^, suggesting Rab33B might serve to recruit the Ric1-Rgp1 complex to the Golgi. However, this region of human RIC1, which lies C-terminal to the GEF domain identified here, is not present in budding yeast Ric1. Therefore another mechanism appears to be responsible for controlling the precise localization of Ric1-Rgp1 in budding yeast, and perhaps in metazoan cells as well, which will be important to investigate in future studies.

A distinguishing feature of all Rab GTPases is the flexible, C-terminal tail referred to as the hypervariable domain (HVD). The HVD consists of a stretch of ∼30 unstructured residues that are conserved among species but divergent between Rabs. The HVD is typically post-translationally modified by the addition of prenyl groups to the C-terminal cysteines which insert into the membrane, flexibly tethering the Rab to the membrane surface upon activation. Given the divergence of HVD sequences between different Rabs, they are capable of providing recognition specificity for their regulators. The HVD has been reported to be important for some, but not all, Rab proteins^61–65^. Structural and functional analysis of the TRAPP complexes revealed that direct interactions with the HVDs of their Rab substrates are important for their substrate specificity^26,36,37,47^. The architecture of the Mon1-Ccz1 GEF complex bound to Rab7^66,67^ suggests this GEF may also directly bind to the HVD of its substrate, although an interaction with the Rab7 HVD has not yet been visualized. For other structurally characterized interactions between Rab GTPases and their GEFs, the constructs used for structural studies did not include the Rab HVD, and often did not include the entire GEF protein. Therefore, the available structural data suggest a common mechanism in which the HVDs of many GTPases contribute to the specificity of their GEFs via direct interactions with these otherwise unstructured regions.

## METHODS

### Plasmid and strain construction

Standard molecular biology techniques were used to construct strains (Table 2) and plasmids (Table 3).

**Table 2:**
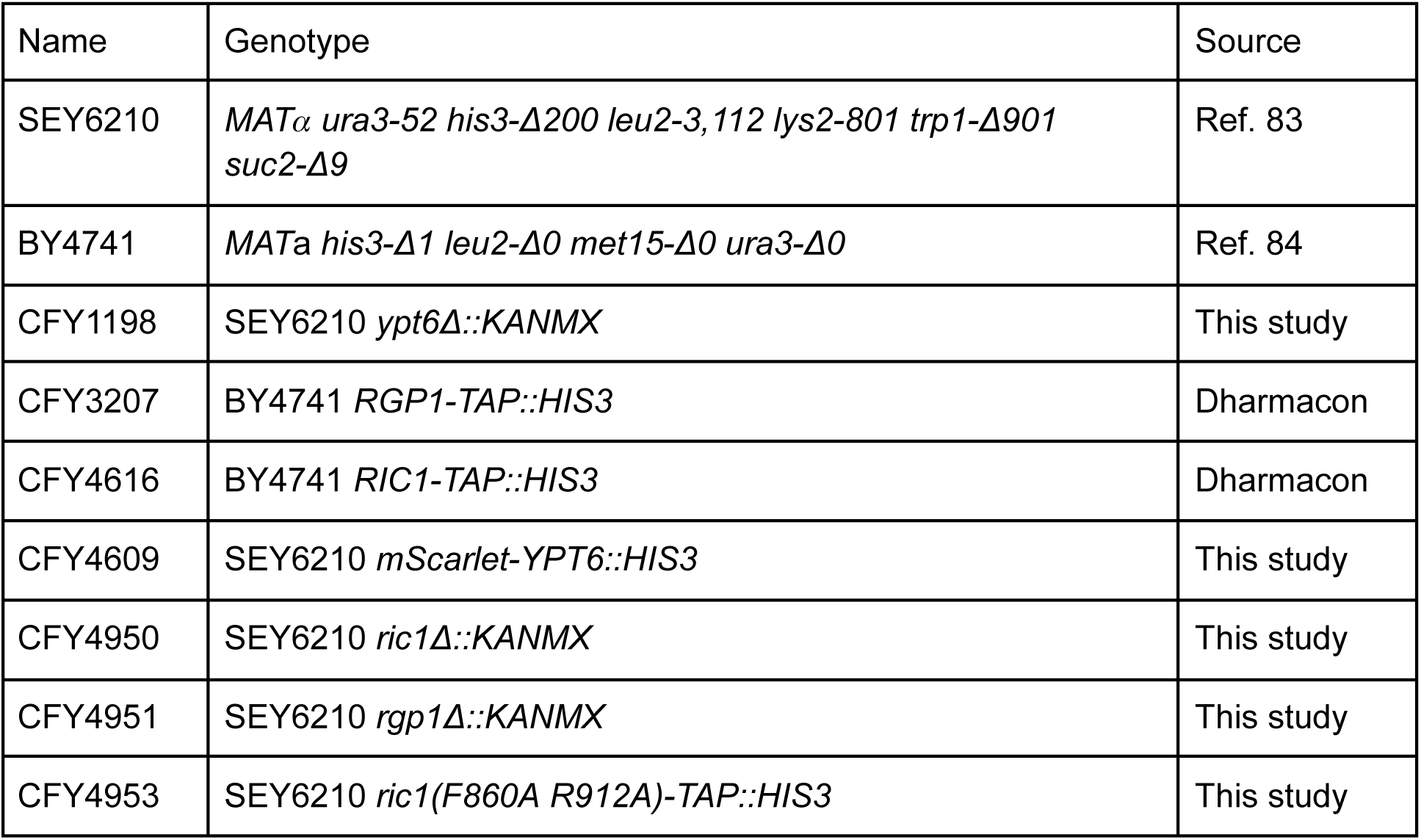
Yeast strains used in this study.

**Table 3:**
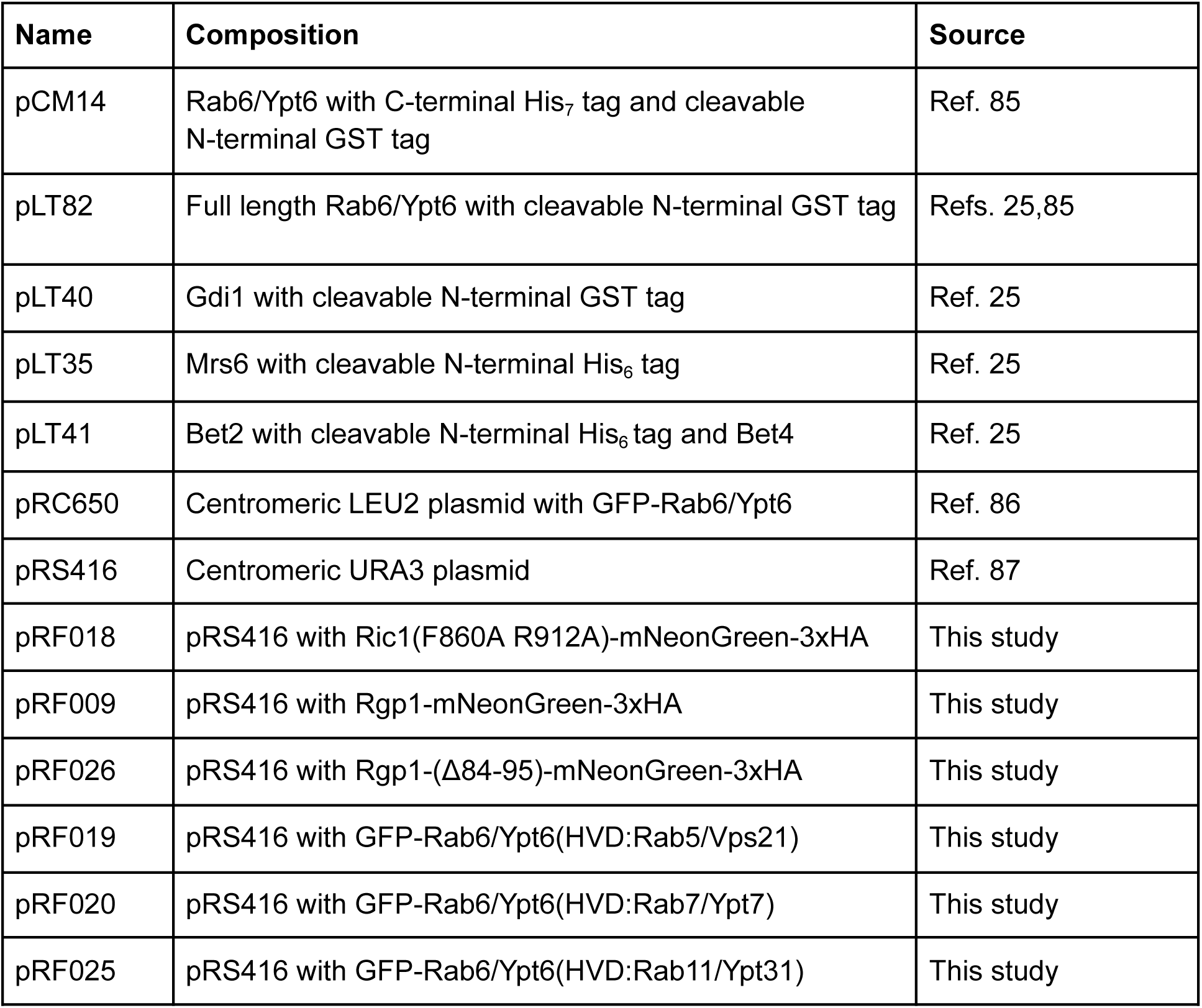
Plasmid reagents used in this study.

| **Name** | **Composition** | **Source** |
| --- | --- | --- |
| pCM14 | Rab6/Ypt6 with C-terminal His<sub>7</sub> tag and cleavable N-terminal GST tag | Ref. 85 |
| pLT82 | Full length Rab6/Ypt6 with cleavable N-terminal GST tag | Refs. 25,85 |
| pLT40 | Gdi1 with cleavable N-terminal GST tag | Ref. 25 |
| pLT35 | Mrs6 with cleavable N-terminal His<sub>6</sub> tag | Ref. 25 |
| pLT41 | Bet2 with cleavable N-terminal His<sub>6</sub> tag and Bet4 | Ref. 25 |
| pRC650 | Centromeric LEU2 plasmid with GFP-Rab6/Ypt6 | Ref. 86 |
| pRS416 | Centromeric URA3 plasmid | Ref. 87 |
| pRF018 | pRS416 with Ric1(F860A R912A)-mNeonGreen-3xHA | This study |
| pRF009 | pRS416 with Rgp1-mNeonGreen-3xHA | This study |
| pRF026 | pRS416 with Rgp1-(Δ84-95)-mNeonGreen-3xHA | This study |
| pRF019 | pRS416 with GFP-Rab6/Ypt6(HVD:Rab5/Vps21) | This study |
| pRF020 | pRS416 with GFP-Rab6/Ypt6(HVD:Rab7/Ypt7) | This study |
| pRF025 | pRS416 with GFP-Rab6/Ypt6(HVD:Rab11/Ypt31) | This study |

### Protein purification

The Ric1-Rgp1 complex was purified using tandem affinity purification from yeast. 12-24 liters of yeast strains with chromosomally tagged Ric1-TAP or Rgp1-TAP were grown at 30°C shaking in YPD and harvested at an O.D. of ∼2. Cell pellets were washed once then resuspended in a 1:1 ratio with CHAPS lysis buffer + protease inhibitors (Roche Complete) and frozen dropwise in liquid nitrogen before freezer-mill lysis. Clarified lysate was incubated with Sepharose 6B resin to remove non-specific binding proteins prior to incubation with IgG resin. Bound IgG resin was transferred to a gravity chromatography column to wash and buffer exchange. For cryoEM samples Rab6 was added in excess together with calf intestine alkaline phosphatase (Invitrogen) to hydrolyze bound GDP. Ric1-Rgp1 or Ric1-Rgp1-Rab6 was cleaved from the resin by overnight incubation with TEV protease at 4°C. Eluate from the IgG resin was collected and subjected to a second affinity purification with calmodulin resin followed a wash and elution in 150ul fractions. Fractions containing protein complex were concentrated then further enriched and buffer exchanged by gel filtration using a Superdex 200 Increase 3.2/300 column. The final buffer contained 25 mM HEPES pH 7.4, 150 mM NaCl, 1 mM dithiothreitol. Pooled fractions were concentrated to 1-2 mg/ml. 12 L of yeast resulted in a typical yield of roughly 20 μg of purified complex.

All recombinant proteins from *E. coli* were freshly transformed and expressed in Rosetta2 cells. Rab6, Gdi1, Mrs6, Bet2, and Bet4 were purified following previously established methods ^25^. Yeast Rab6 and Gdi1 tagged with GST were purified with glutathione affinity resin and eluted by cleavage with PreScission protease to produce an untagged final product. 6xHis-tagged Mrs6 and 6xHis-Bet2-Bet4 were purified following standard procedures with nickel affinity resin. All recombinant proteins were aliquoted and flash frozen in liquid nitrogen.

### CryoEM sample preparation

3 μl of sample from freshly concentrated gel filtration fractions was applied to glow-discharged Quantifoil or UltrAufoil R1.2/1.3 300-mesh gold-support grids. For datasets imaged with detergent, octyl-□-glucoside was added shortly before applying sample to a final concentration of 0.07%-0.1%. Grids were blotted for 2-5 seconds with blot force 0 at 4°C and 100% humidity and immediately plunge-frozen in liquid ethane using a FEI Mark IV Vitrobot.

### CryoEM data collection, processing, and model building

CryoEM data collection was performed using a Thermo Fisher Scientific Talos Arctica operated at 200 keV equipped with a Gatan K3 detector operated in counting mode with 0.5X-binning (super-resolution mode) and a Gatan BioQuantum energy filter with a slit width of 20 eV.

Microscope alignments were performed on a gold diffraction cross-grating following published procedures for a 2-condenser instrument^68,69^. Parallel conditions were determined in diffraction mode at the imaging magnification. A 50 μm C2 aperture size was chosen and the spot size was set so that the dose rate at the detector was ∼25 e-/physical pixel/sec over vacuum. A 100 μm objective aperture was used during data collection.

SerialEM ^70^ was used for automated data collection of six independent datasets of 50 frame fractionated exposures with a total dose of ∼50 e-/Å^2^ so that the dose per frame was ∼1 e-/Å^2^. Data were collected at defocus values ranging from -0.8um to -1.5um with a nominal magnification of 63,000x resulting in a physical pixel size of 1.25 Å. Preliminary screening identified a strong preferred orientation bias that precluded 3D reconstruction of an interpretable map. We therefore collected data from both flat and tilted grids. A total of 2931 exposures were used for data processing, with 2142 of those exposures obtained from grids tilted to 30°.

The cryoEM data processing tools RELION^71,72^, MotionCor2^52,73^, GCTF^52,73,74^, and CryoSPARC^75^ were used to correct beam-induced motion, estimate CTF parameters, pick and sort particles, and to perform refinements, classifications, and reconstructions. An *ab inito* reference model for Ric1-Rgp1-Rab6 was generated with cryoSPARC by combining select 2D class averages from flat and tilted datasets. For tilted datasets, per particle defocus estimates were refined using local GCTF. All refinement and classification steps were carried out in RELION. Initial high-resolution refinements exhibited Rab6 density only at a low threshold.

Signal subtraction and fixed angle 3D classification were used to further isolate particles of the Rab-bound complex. The final refinement resulted in a reconstruction with a resolution of 3.3 Å (FSC 0.143 cut-off). Local-resolution filtering^76^ was used to generate the map used for model building and figures.

The initial atomic model for Ric1-Rgp1-Rab6 was built *de novo* using ‘Map-to-Model’^77^ in Phenix^78^. The output from map to model was rebuilt manually in Coot^79^. Model refinement and validation was performed using Phenix real-space refinement^80^ and Phenix cryoEM validation^81,82^.

### Liposome preparation

Liposomes were prepared with a membrane composition similar to the lipid profile observed at late secretory pathway and TGN derived vesicles (Table 4)^27^. Lipids solubilized in chloroform were mixed and dried under a stream of argon or nitrogen. Unilamellar vesicles were prepared by hydrating the dried film in HK buffer (25 mM HEPES pH 7.4, 150 mM KOAc, 1 mM DTT) at 37°C with occasional mixing for 1 hour followed by 2 minutes of sonication in a bath sonicator then 3-5 freeze thaw cycles. As a final step, the liposomes were extruded for a total of 15 passes through a 100nm filter (Whatman).

**Table 4:**
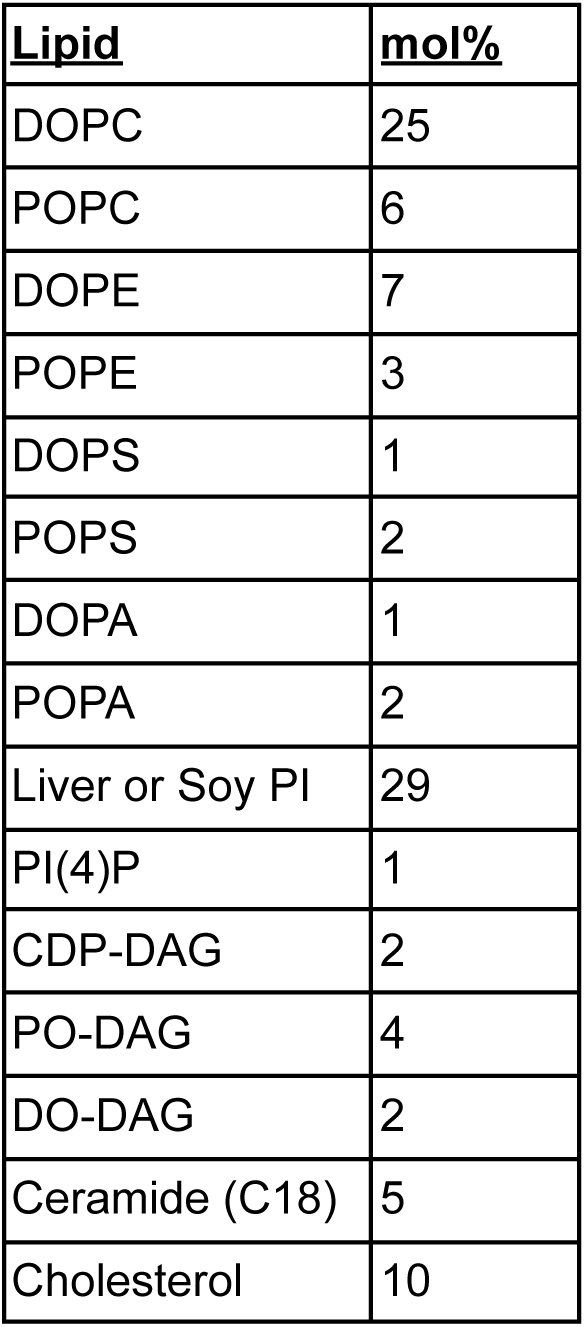
TGN lipid mix.

| <u>Lipid</u> | <u>mol%</u> |
| --- | --- |
| DOPC | 25 |
| POPC | 6 |
| DOPE | 7 |
| POPE | 3 |
| DOPS | 1 |
| POPS | 2 |
| DOPA | 1 |
| POPA | 2 |
| Liver or Soy PI | 29 |
| PI(4)P | 1 |
| CDP-DAG | 2 |
| PO-DAG | 4 |
| DO-DAG | 2 |
| Ceramide (C18) | 5 |
| Cholesterol | 10 |

### Prenylated Rab6-mantGDP-Gdi1 preparation

The prenylated Rab6-GDI complex bound to fluorescent mant-GDP was prepared as previously described^25^. Recombinant Rab6 was first loaded with the fluorescent nucleotide analog mantGDP. A 1ml exchange reaction was prepared by combining purified full length Rab6 at a final concentration of 40 μM with 200 μM mantGDP and 20mM EDTA in prenylation buffer (20 mM HEPES, 150 mM NaCl, 2 mM MgCl_2_, 1 mM DTT) and incubated for 30 min at 30°C. Exchange was completed by adding MgCl_2_ to a final concentration of 25 mM. Excess mantGDP and MgCl_2_ was removed by buffer exchange with a 5 mL Zeba spin desalting column (ThermoFisher) following standard procedures. The C-terminal cysteines of Rab6 were prenylated by combining Rab6-mantGDP, Gdi1, 6xHis-Bet2-Bet4, and 6xHis-Mrs6 in a 10:10:1:1 ratio in prenylation buffer containing a final concentration of 120 μM geranyl-geranyl pyrophosphate and 25 μM mantGDP. After allowing the reaction to proceed for 1 hour at 37°C, Ni-NTA resin was added to remove the tagged geranylgeranylation machinery proteins. Stoichiometric prenylated Rab6-mantGDP in complex with Gdi1 was subsequently buffer exchanged and isolated by gel filtration.

### GEF activity assay

Rab6 nucleotide exchange was monitored using an established physiological *in vitro* RabGEF assay^25,26^. MantGDP exhibits increased fluorescence (365-nm excitation/440-nm emission) when bound to Rab GTPases. Exchange reactions were prepared in a total volume of 150 μl with HKM buffer (20mM HEPES pH 7.4, 150 mM KoAc, 2 mM MgCl_2_, 1 mM DTT) by first combining all of the components except the GEF in a quartz cuvette at 30°C. The reaction was allowed to equilibrate for four minutes before adding wild type or mutant Ric1-Rgp1, and measuring fluorescence change over time. The reaction concentrations of each component were 333 μM lipids, 200 μM GTP, 1 μM Rab6-mantGDP-Gdi1, and 250 nM Ric1-Rgp1 complex.

### Antibodies

For immunoblots, the following antibodies were used at the following concentrations: mouse anti-HA (Roche 12CA5), 1:500; rabbit anti-TAP (Invitrogen CAB1001), 1:1000; mouse anti-GFP (Santa Cruz sc-9996), 1:500; HRP-conjugated anti-rabbit (Cytiva NA934V) 1:10,000; HRP-conjugated anti-mouse (Cytiva NXA931V), 1:10,000.

### Fluorescence microscopy

Cells were grown to early log phase at 30°C in synthetic drop out media. All images were collected with a DeltaVision RT widefield deconvolution microscope using a 100x 1.35 N.A. oil immersion objective lens. Acquisition and deconvolution were performed in the SoftWoRx software.

### Image analysis

Deconvolved images were preprocessed in FIJI to generate 200 pixel cropped unnormalized 16-bit image stacks containing 1-2 budding cells using a custom plugin (https://github.com/ryanfeathers/Stack_Box.git). For visualization all histograms were normalized to the same intensity range for each channel and inverted in FIJI. Colocalization analysis was performed in CellProfiler 4.2.1. A single z-slice from each cropped stack was first subjected to subtraction of the median raw intensity value to reduce background followed by Manders’ correlation coefficient measurement.

### Statistical analysis

Statistical significance was calculated in R using an unpaired two-tailed t-test with Welch’s correction. For box and whisker plots the median is denoted by a line, and the box extends from the first quartile to the third quartile of the data. The whiskers extend from the box to the farthest data point lying within 1.5x of the inter-quartile range from the box. All individual data points are also shown.

### Data Deposition

CryoEM maps have been deposited in the EMDB (EMD-43997) and structure coordinates have been deposited in the RCSB PDB (9AYR).

## ACKNOWLEDGEMENTS

We thank L. Kourkoutis for advice and assistance regarding instrumentation and data collection. We acknowledge the Cornell Center for Materials Research (CCMR), especially K. Spoth and M. Silvestry-Ramos, for access and support of electron microscopy sample preparation and data collection. We thank R. Collins for the gift of a GFP-Rab6/Ypt6 plasmid. We thank members of the Fromme lab for helpful advice and discussions. This study was funded by NIH grant R35GM136258 to J.C.F. and a Ford Foundation Predoctoral Fellowship to J.R.F. The CCMR is supported by NSF grant DMR-1719875.

## AUTHOR CONTRIBUTIONS

J.R.F. and J.C.F. conceived the study and experimental design. J.R.F. and R.C.V performed experiments. J.R.F, R.C.V, and J.C.F analyzed data and made figures. J.C.F. supervised the project and obtained funding. J.R.F. and J.C.F. wrote the manuscript and R.C.V. edited the manuscript.

## SUPPLEMENTARY INFORMATION

**Supplementary Figure S1:**
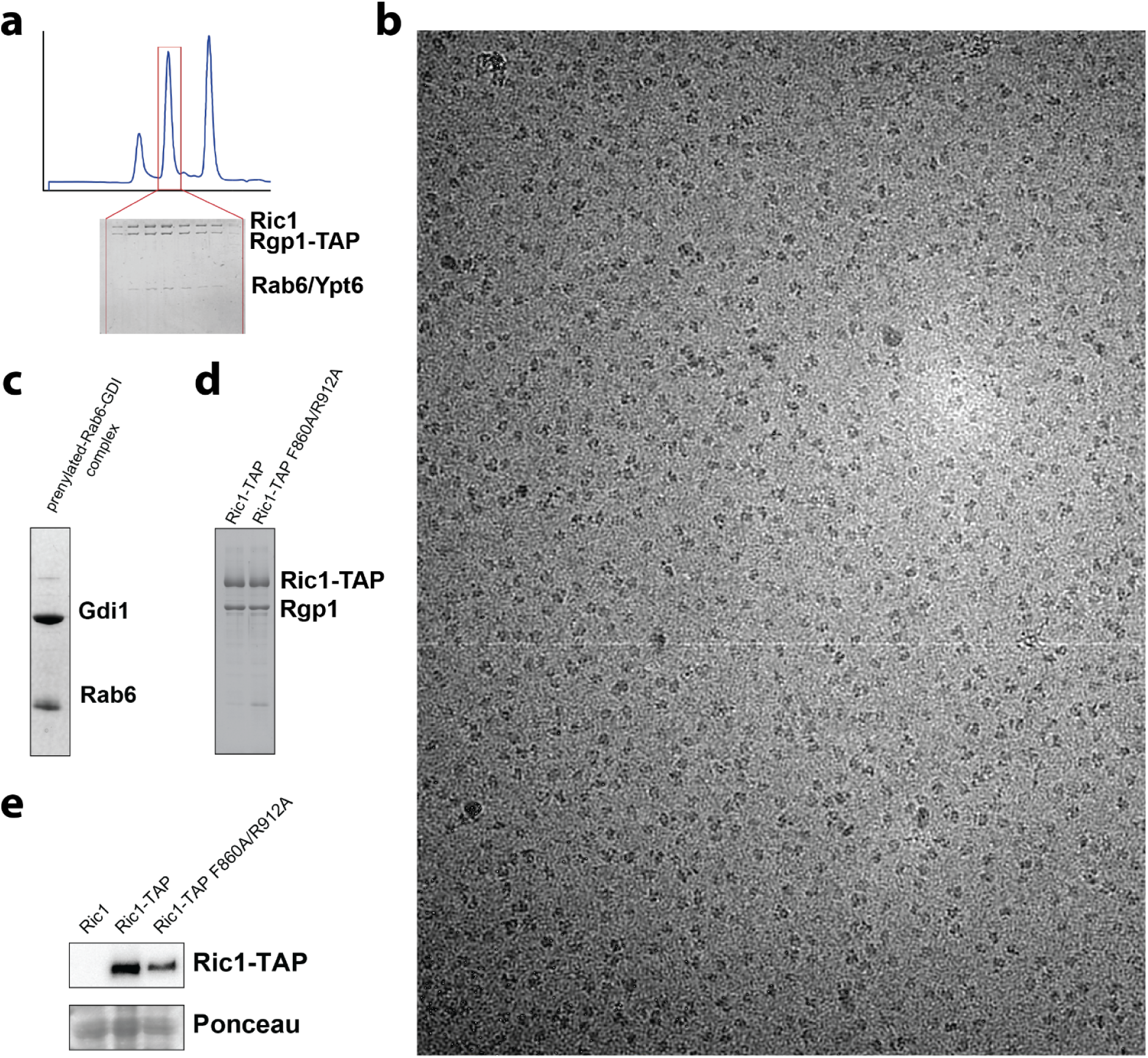
Complex preparation and cryoEM imaging data. a) Gel-filtration chromatography trace and corresponding coomassie-stained SDS-PAGE gel for the final purification step of the Ric1-Rgp1-Rab6 complex. b) Example cryoEM micrograph of the purified complex. c) Coomassie-stained SDS-PAGE gel of the purified prenyl-Rab6-GDI complex used as a substrate for the *in vitro* GEF assays. d) Coomassie-stained SDS-PAGE gel of the purified WT and mutant Ric1-Rgp1 complexes used for the *in vitro* GEF assays. e) Immunoblot to determine the protein levels of WT and mutant endogenously tagged Ric1-TAP proteins.

**Supplementary Figure S2:**
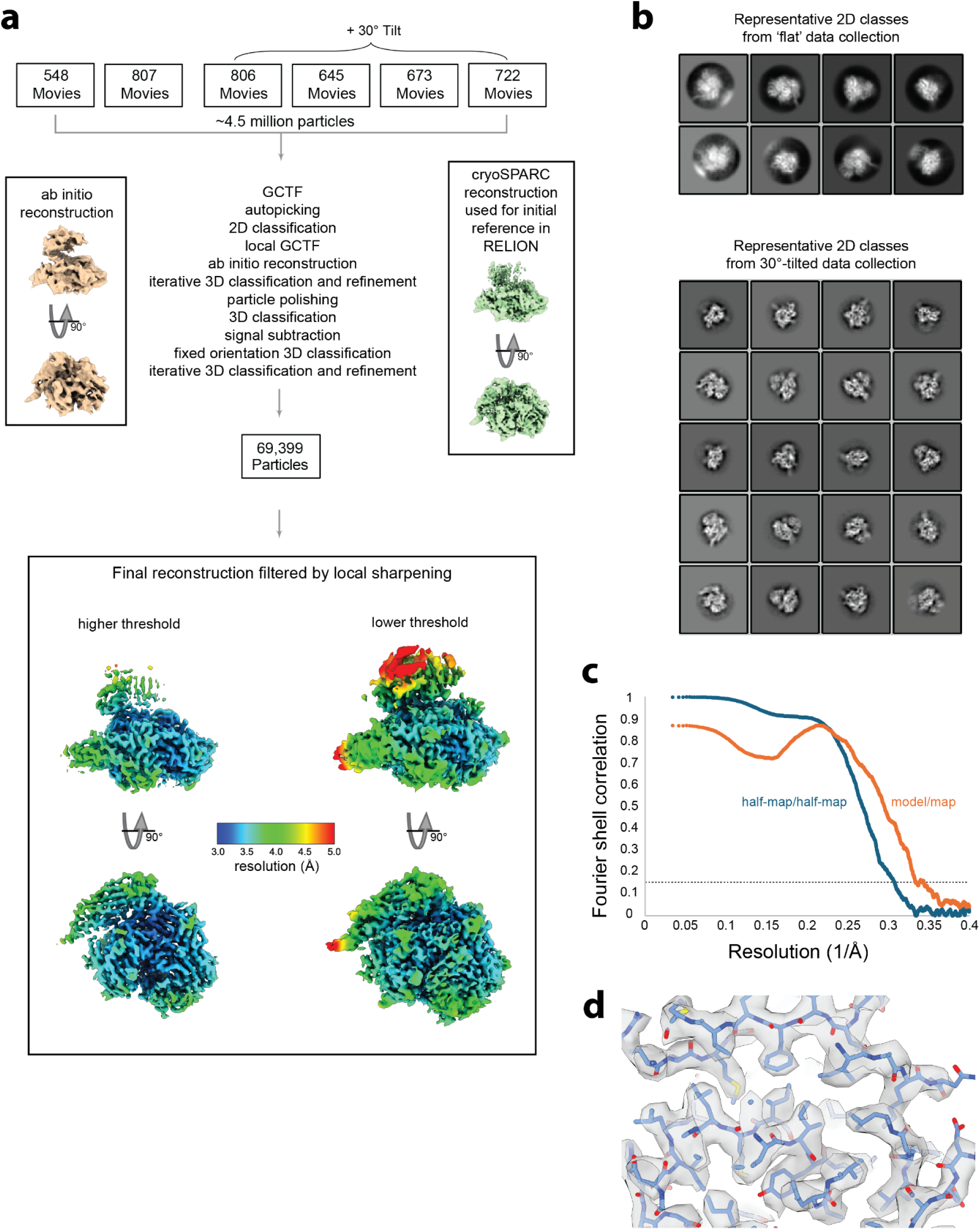
CryoEM data processing, refinement, and reconstruction. a) CryoEM data collection and processing workflow, including images of the 3D reconstructions at different stages of data processing. b) Example 2D class averages calculated from data collected on flat and tilted grids. c) FSC curves for half-map/half-map from the final refinement and model/map of the final atomic model. The dotted line indicates the 0.143 FSC cutoff. d) View of the sharpened cryoEM density superimposed on the final atomic model.

**Supplementary Figure S3:**
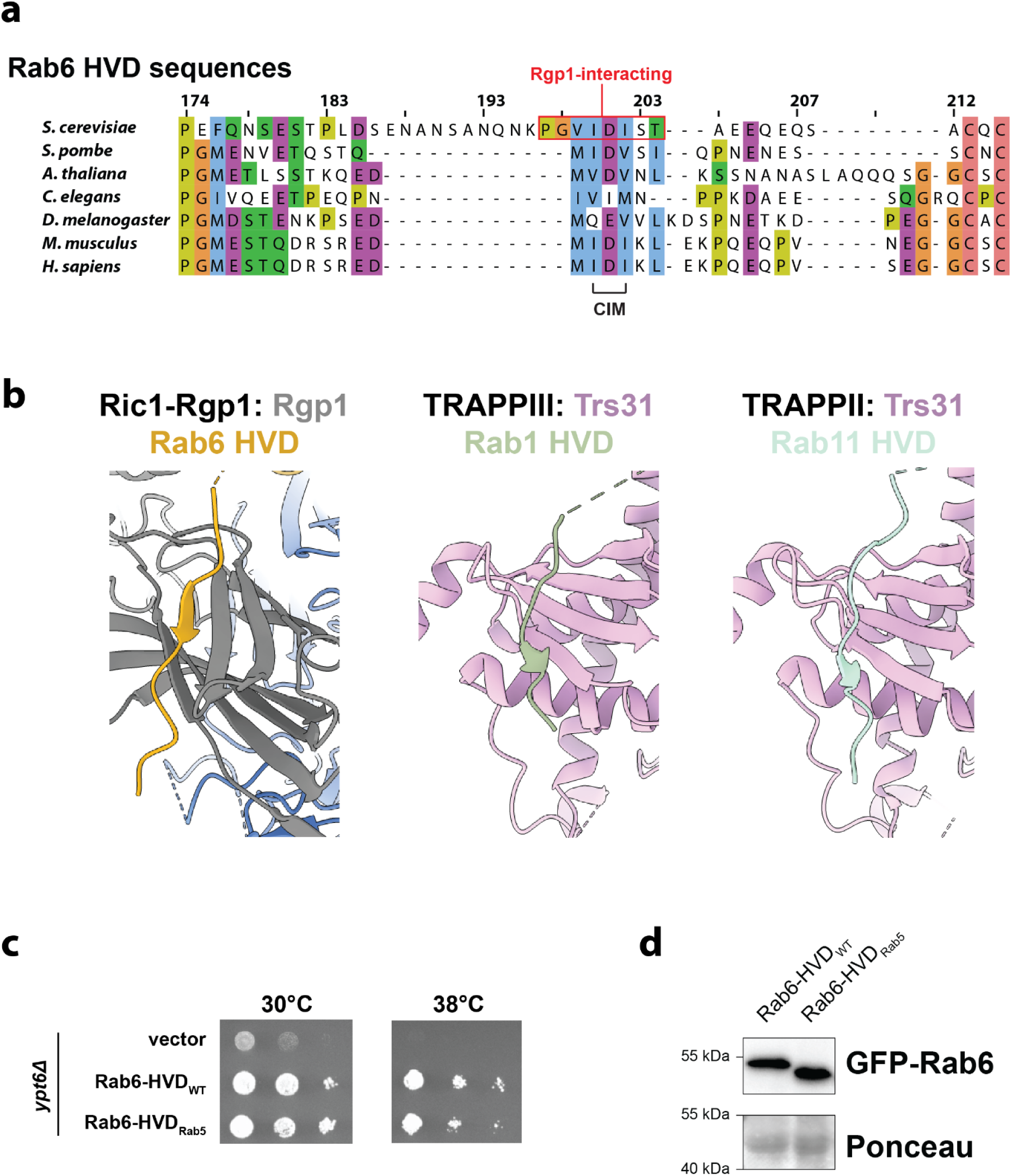
Binding to the HVD may be a general feature of RabGEFs. a) Sequence alignment of the HVDs from several Rab6 paralogs. The ‘CIM’ motif known to bind to the geranyl-geranyl transferase machinery is denoted, and the residues of S. cerevisiae Rab6 that interact with Rgp1 in the cryoEM structure are outlined with a red box. b) Comparison illustrating how the Trs31 subunit of the TRAPPII and TRAPPIII complexes binds to the HVD of Rab11 and Rab1, respectively (PDB: 7U05 and 7KMT)^36–38^. c) Rab6 complementation test. d) Immunoblot to determine the protein levels of WT and chimeric mutant GFP-tagged Rab6/Ypt6 proteins expressed as an extra copy from a centromeric plasmid in yeast cells.

**Supplementary Figure S4:**
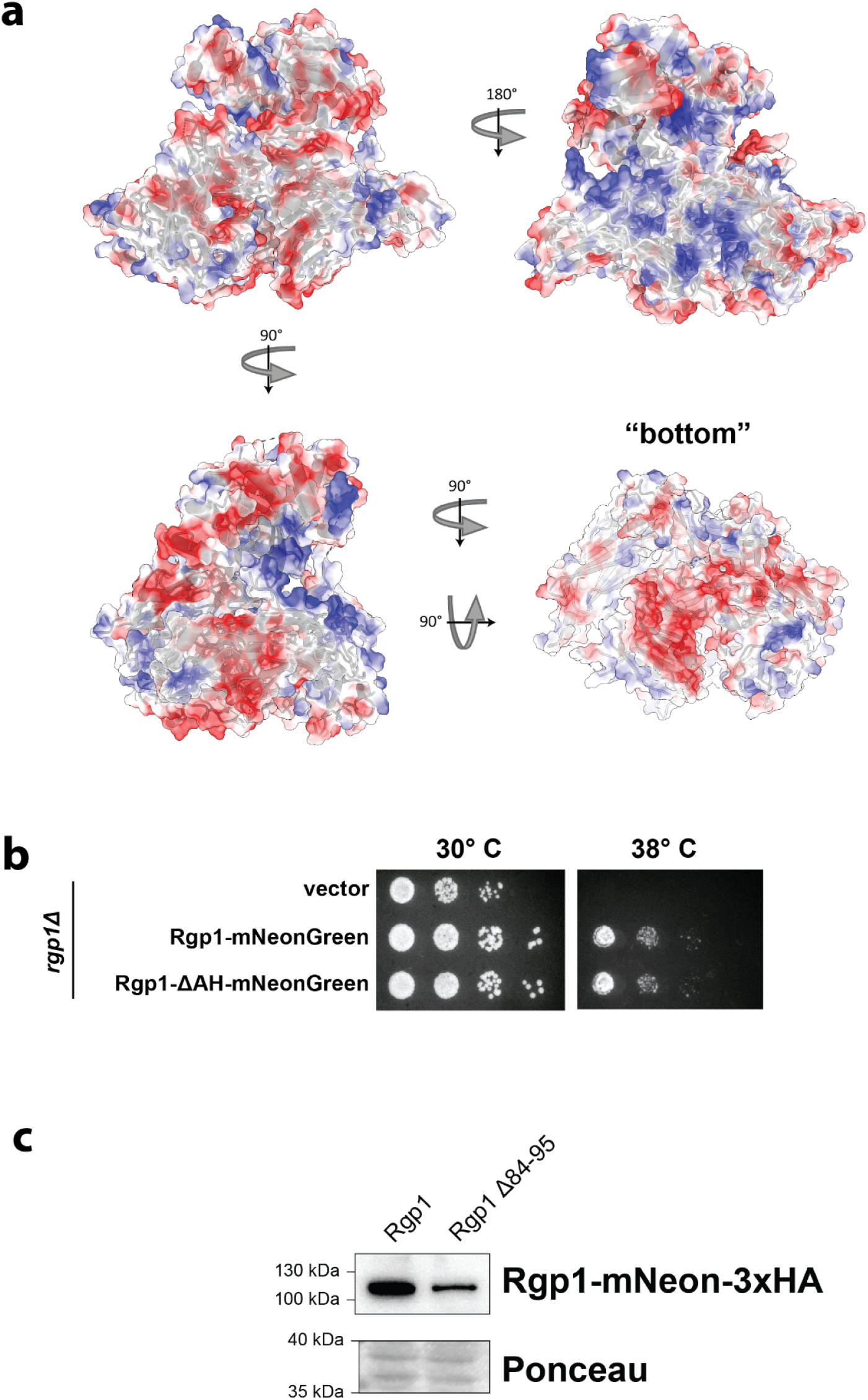
Analysis of Ric1-Rgp1 surface electrostatics. a) Electrostatic surface representation of the Ric1-Rgp1-Rab6 atomic model determined by cryoEM. The “bottom” surface which we predict faces the membrane is indicated. b) Complementation test (cells lacking Ric1-Rgp1 function are temperature sensitive). c) Immunoblot to determine the protein levels of WT and mutant mNeonGreen-3xHA-tagged Rgp1 proteins expressed on a centromeric plasmid in *rgp1Δ* yeast cells.

**Video S1: Rab6 activation by the Ric1-Rgp1 complex**

Initially, Rab6, colored in yellow, is shown bound to GDP, colored in red, representing the GTPase in the inactive state. Rab6 is shown diffusing in and "morphing" to the nucleotide free conformation upon association with the GEF domain of Ric1 colored in blue. In this conformation, the GDP freely diffuses and is exchanged for GTP which is colored in green. Rab6 undergoes a conformational change once again and is shown transitioning to the GTP bound state. The GTP-bound Rab6 conformation is not compatible with stable binding to the Ric1-GEF domain, so activated Rab6 dissociates from the GEF. This video was made in ChimeraX^88^ using models generated in SWISS-MODEL^89^ based on PDB 2FE4 (human Rab6B-GDP)^90^, PDB 5LEF (human RAB6A-GTP)^91^, and PDB 9AYR (the Ric1-Rgp1-Rab6 structure reported in this work).

